# An IGF2-anchored oncofetal state and relapse-associated transcriptomic module define systemic progression risk in early-onset colorectal cancer

**DOI:** 10.64898/2026.05.18.725955

**Authors:** Elena Licitra-Rosa, Giulia Mantini, Federica Persiani, Eleonora Ponterio, Sebastiano Di Bella, Laura Lorenzon, Giulia Scaglione, Miriam Caimano, Domenico D’Ugo, Lisa Salvatore, Maria Alessandra Calegari, Gianfranco Zannoni, Giorgio Stassi, Ruggero De Maria

## Abstract

The epidemiological surge of early-onset colorectal cancer (EOCRC) is characterized by accelerated biological kinetics and disproportionately high rates of systemic relapse following curative-intent surgery. Because standard anatomical staging (TNM) lacks the resolution to accurately capture the intrinsic regenerative capacity of microscopic residual disease, we investigated the transcriptomic architecture of post-surgical failure in a strictly defined, curative-intent clinical pan-cohort. Unbiased transcriptomic profiling of the localized (M0) discovery sub-cohort identified IGF2 as the most significantly upregulated correlate of metachronous relapse. High-resolution isoform analysis revealed that this transcriptional output is predominantly driven by the embryonic (P4) and placental (P5) promoters. Systematic allele-specific expression (ASE) analysis supported widespread biallelic IGF2 expression consistent with relaxation of imprinting-domain control. This signal was not restricted to relapsing tumors, suggesting a recurrence-independent oncofetal baseline across the EOCRC spectrum. Because this foundational epigenetic unlocking is functionally insufficient on its own to execute systemic metastasis, we distilled the additional transcriptional plasticity required for dissemination into an internally derived and bootstrap-stabilized 5-gene recurrence-risk module Multivariable analysis across the combined pan-cohort supported an independent association between the high-risk module and systemic relapse (p < 0.001), capturing prognostic dimensions completely unresolved by classical pathological covariates and baseline staging. Ultimately, our findings reframe EOCRC aggressiveness as the product of a dual-hit architecture. This framework resolves the clinical paradox of widespread IGF2 LOI co-existing with heterogeneous outcomes, offering a biologically grounded basis for molecular risk stratification beyond anatomical boundaries.

## Introduction

The epidemiological acceleration of Colorectal Cancer (CRC) incidence among younger demographics has fractured traditional paradigms of pathogenesis and clinical management [1]. Historically relegated to the periphery of oncological focus due to standard age-based screening cut-offs, Early-Onset CRC (EO CRC) is now recognized not merely as a demographic anomaly, but as a biologically distinct clinical and molecular setting [2]. However, current prognostic and interpretative frameworks remain deeply anchored in the mutational and microenvironmental landscapes of late-onset, chronologically aged cohorts [3, 4]. Consequently, these legacy models fail to capture the unique ontogenetic complexity and the intrinsic developmental drivers that fuel the aggressiveness of EO CRC [5].

While EO CRC is uniquely enriched for hereditary cancer syndromes and microsatellite instability (MSI), the great majority of emerging cases are strictly sporadic [6,7]. Paradoxically, these sporadic tumors often exhibit a somatic mutational landscape that is substantially superimposable onto that of older cohorts, barring minor deviations [8,9]. Although the modern exposome is undeniably fueling the epidemiological surge, it fails to fully explain the accelerated biological kinetics of the disease [10, 11]. This discrepancy imposes a profound mechanistic challenge, effectively fracturing the deterministic rigidity of the classic Vogelstein adenoma-carcinoma sequence, which relies on a strictly time-dependent accumulation of genetic hits [12].

This mechanistic impasse suggests that the ‘energy’ required to drive malignant transformation in younger patients operates on a fundamentally different physical and biological level. In average-onset cohorts, tumorigenesis dissipates energy to create chaos within a worn-out, highly entropic tissue environment, while the disruptive kinetics of EO CRC must carve its path through a more structurally intact genomic and cellular landscape [13, 14]. This proposition is actively supported by emerging multi-omics profiles, which demonstrate that EO CRC operates via a distinct molecular architecture characterized by profound DNA methylation perturbations and DNA repair dysregulation, distancing itself from the classical progression models of older cohorts [15].

Crucially, this distinctiveness is not merely genomic, but deeply structural and biophysical. Recent biomechanical phenotyping reveals that EO CRC is marked by early and widespread fibrotic remodeling [16]. Compared to late-onset disease, early-onset tumors and their adjacent normal tissues exhibit significantly increased matrix stiffness, elevated viscosity, and a highly aligned, dense collagen microstructure that mechanically forces epithelial activation and tumor proliferation through enhanced YAP-mediated mechanotransduction.This unique remodeling of the extracellular matrix is further evidenced by disease-specific alterations in structural and inflammatory modulators [17].

Perhaps most strikingly, to overcome the rigid biomechanical barrier of a healthy, non-senescent host environment, EO-CRC appears to hijack primordial networks, exhibiting distinctive “placental-like” biological features [18]. High-resolution proteomics and transcriptomics have revealed that EO-CRC tumors are significantly enriched in proteins and pathways typically confined to early developmental stages, specifically mirroring the embryonic trophoblast. Unable to rely on the entropic degradation of the aging colon, early-onset tumors mimic the invasive strategies of the embryological trophoblast to breach the surrounding tissues.

Clinically, this divergent biology translates into a highly aggressive pattern of systemic escape.This reality is reflected both in the disproportionately higher proportion of patients presenting with de novo metastatic (M1) disease [19], and in the failure of local interventions. Even when intercepted at a strictly localized (M0) stage and managed with curative-intent surgery, younger patients endure significantly higher rates of metachronous relapse compared to traditional, late-onset cohorts [20].

To successfully execute this peripheral dissemination, disseminated tumor cells must not only exploit the immune-privileged, highly vascularized placental blueprint to evade clearance, but also fundamentally rewire their cellular energetic programs to maximize nutrient extraction from the surrounding microenvironment [21,22].

Evolutionarily, this metabolic and developmental reprogramming is centrally governed by paternally expressed imprinted genes that are specifically designed to demand and extract maternal resources, maximizing fetal growth, vascularization, and placental expansion [23-25]. By hijacking these mechanisms, tumor cells can aggressively sequester host nutrients, providing the energetic supply required to fuel their rapid kinetics [22].

While this developmental reversion secures the energetic capital for local expansion, the phenotypic fluidity required for systemic dissemination and survival in a distant organ necessitates a distinct layer of real-time transcriptomic adaptability. To identify the specific transcriptomic machinery required to stabilize this archaic engine and secure successful peripheral dissemination, we designed a step-wise transcriptomic investigation. We initially explored public TCGA-COADREAD cohorts to map the foundational oncofetal baseline of EO-CRC. Subsequently, to characterize the molecular architecture of systemic failure, we transitioned to an internal, curative-intent clinical cohort of EO-CRC patients (n=96), comprising localized (M0, n=77) and fully resected, macroscopically disease-free, de novo metastatic (M1, n=19) cases. Through transcriptomic profiling of the localized (M0) sub-cohort, we isolated the specific drivers of metachronous relapse, integrating these vulnerabilities into a derived multi-class prognostic signature. We first validated this hardwired ‘executor’ module in the M1-resected extreme, and subsequently evaluated it across the combined curative-intent pan-cohort, ultimately demonstrating its capacity to stratify the risk of systemic relapse independently of classical clinical staging.

## Results

### Exploratory transcriptomic profiling reveals an immune-privileged, oncofetal baseline in genomically stable EO-CRC

To establish the baseline molecular architecture of early-onset disease, we first conducted an exploratory transcriptomic analysis utilizing public COAD/READ cohorts from the TCGA. To strictly eliminate confounding transcriptomic noise derived from hypermutated phenotypes, systemic therapies, and variant histologies (e.g., mucinous or signet-ring cell carcinomas), we rigorously restricted our analysis to treatment-naive, genomically stable (MSS, POLE-wildtype) classical adenocarcinomas (Not Otherwise Specified; NOS). Differential gene expression was evaluated by contrasting the early-onset group (<50 years, n=39) against both average-onset (50–70 years, n=144) and late-onset (> 70 years, n=129) references [Fig. 1A, 1B].

**Figure 1.**
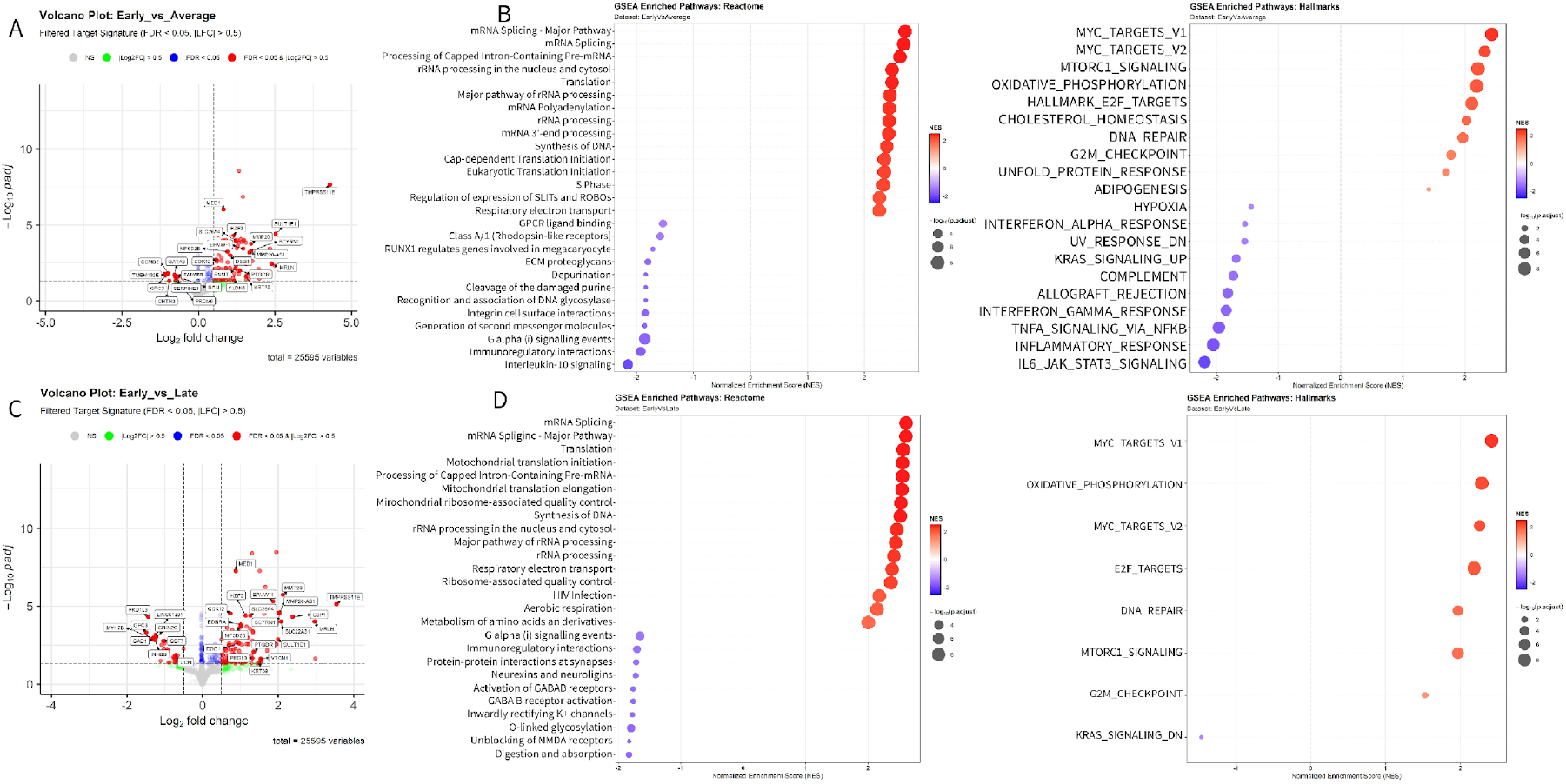
Transcriptomic characterization and functional enrichment of early-onset colorectal adenocarcinomas (COAD/READ). **(A)** Volcano plot showing differential gene expression (DGE) analysis between the early-onset (< 50 years, n=39) and average-onset (50–70 years, n=144) cohorts. Statistically significant genes (FDR < 0.05, |Log2FC| > 0.5) are highlighted in red, with top-ranking genes annotated. **(B)** Dot plots of Gene Set Enrichment Analysis (GSEA) results for the Early vs. Average comparison, utilizing Reactome (left) and Hallmark (right) ontologies. Pathways are ordered by Normalized Enrichment Score (NES). A significant enrichment (positive NES, red) in RNA splicing, translation processes, and MYC targets is observed in early-onset tumors, alongside a downregulation (negative NES, purple) of inflammatory and interferon-mediated signaling signatures. **(C)** Volcano plot depicting DGE analysis for the comparison between early-onset and late-onset (> 70 years, n=129) cohorts. Significance thresholds remain unchanged (FDR < 0.05, |Log2FC| > 0.5). **(D)** GSEA results for the Early vs. Late comparison. Reactome and Hallmark ontologies confirm the sustained activation of RNA metabolism and oxidative phosphorylation, along with the upregulation of MYC and E2F signaling in the early-onset group compared to older patients.

This bioinformatic distillation revealed a highly coordinated upregulation of distinct oncofetal, structural, and metabolic networks. Remarkably, this transcriptomic signature was highly conserved irrespective of the chronological age of the comparator group, confirming EO-CRC as a distinct biological entity rather than a relative demographic extreme.

At the microenvironmental level, this distinct biological identity translates into a profound structural remodeling of the extracellular niche. We observed the extreme upregulation of matrix-degrading enzymes, including *TMPRSS11E* and *MMP20*, a transcriptomic profile consistent with the active biomechanical breakdown of the dense, highly aligned early-onset extracellular matrix [16]. Operating in tandem with this physical remodeling, we observed a massive overexpression of *SULT1E1*, an estrogen sulfotransferase, suggesting the localized modulation of the estrogenic microenvironment.

Operating within this structurally and hormonally secured microenvironment is the targeted derepression of early developmental networks, driving an active ontogenetic reversion. Specifically, we identified the coordinated reactivation of a placental immune-evasion trinity: *ERVW-1* (Syncytin-1), the paternally imprinted factor *PEG10*, and the canonical placental T-cell checkpoint *VTCN1* (B7-H4), supporting the hypothesis that EO-CRC intrinsically co-opts a strategy of trophoblast-like mimicry [26-28]. Furthermore, this developmental state-shift was characterized by a paradoxical transdifferentiation, evidenced by the significant upregulation of the stratified squamous structural components *DSG1* and *KRT39*.

Providing the foundational epigenetic framework to orchestrate and sustain this massive transcriptomic plasticity was the highly significant upregulation of master transcriptional regulators, notably *MED1*, consistent with global enhancer reprogramming [29,30], and *CDK12*, whose overexpression is consistent with the aggressive transcriptional elongation required to sustain large, newly activated oncofetal loci [31]. To capture the overarching functional state driving this phenotype, we performed Gene Set Enrichment Analysis (GSEA) across Reactome, Hallmark, KEGG, and GO ontologies [Fig. 1C, 1D; Supplementary Fig. S1, S2]. Consistent with *CDK12*-mediated transcriptional elongation, EO-CRC exhibited a broad enrichment in pathways governing RNA processing—such as mRNA splicing and rRNA/pre-mRNA maturation—alongside the profound upregulation of adaptive metabolic and stress-response networks, highlighted by MYC and mTORC1 signaling, oxidative phosphorylation, and the Unfolded Protein Response (UPR). Conversely, EO-CRC tumors demonstrated a marked negative enrichment for canonical inflammatory and immune-activating cascades (including *IFN-γ, TNF-α*, and *IL6/JAK/STAT3* signaling) compared to later-onset cohorts, which remained significantly enriched for vestigial colonic physiological functions such as digestion and absorption.

Together, these TCGA data depict genomically stable EO-CRC not merely as a chronologically accelerated tumor, but as a highly plastic entity that exploits both archaic and divergent programs to neutralize the aggressive immune competency of a younger host, engineering a highly shielded, pseudo-placental invasive niche. However, while this global oncofetal baseline maps the intrinsic biology of the disease, the defining clinical challenge of EO-CRC lies in its profound capacity to overcome maximal therapeutic interventions. The ultimate threat is the failure of surgical curative intent—where microscopic residual clones successfully execute a systemic relapse even after the aggressive resection of all macroscopic disease, an approach increasingly deployed even in metastatic scenarios to maximize long-term survival. To decode the specific transcriptomic machinery driving this post-surgical regeneration, we deployed our internal curative-intent clinical cohort (n=96, Supplementary Table 1). We structured our primary discovery phase exclusively within the localized (M0) sub-cohort (n=77), directly contrasting patients who subsequently suffered a surgical failure (metachronous relapse) against disease-free comparators.

### Transcriptomic profiling identifies the *IGF2* locus as the primary correlate of EOCRC recurrence

To delineate the transcriptomic landscape driving clinical recurrence in early-onset disease, we performed an unbiased differential gene expression analysis contrasting localized (M0) patients who subsequently relapsed against disease-free comparators. Global transcriptome analysis identified *IGF2* (Insulin-like Growth factor 2) as the most significantly upregulated locus associated with recurrence. Remarkably, *IGF2* presented not only an extreme log2 fold-change and statistical significance (padj <0.0001) [Fig. 2A], but also maintained a remarkably high absolute baseline expression across the entire cohort. Given the magnitude of this single-locus deregulation, we sought to determine if this deviation was sufficient to orchestrate a global phenotypic shift before dissecting its underlying molecular.

**Figure 2.**
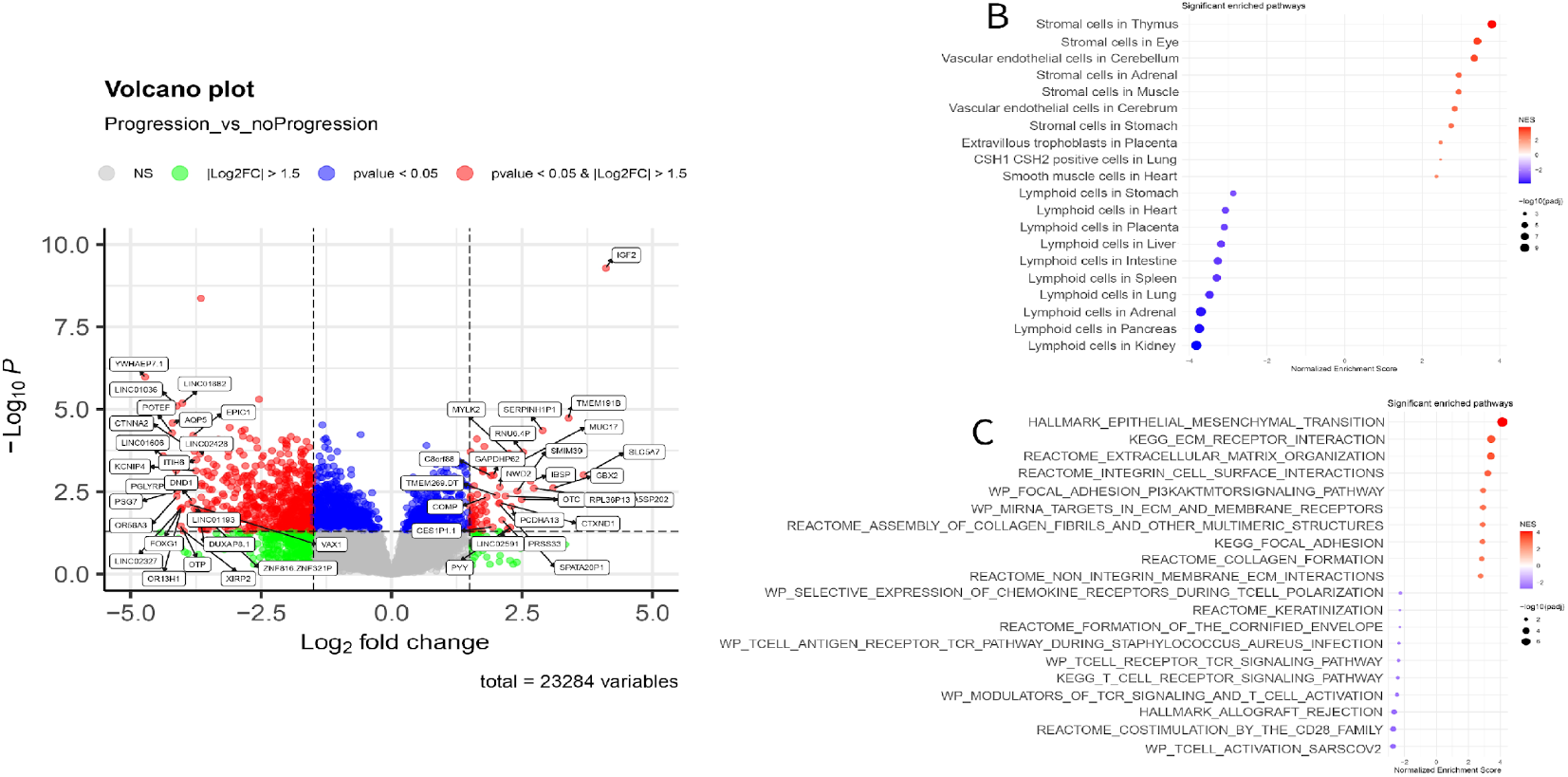
Transcriptomic drivers of M0 patients and microenvironmental remodeling in early-onset disease. (**A**) Volcano plot illustrating differential gene expression associated with disease progression. *IGF2* emerges as the most significantly upregulated transcript in the relapsing cohort. (**B)** GSEA dot plot utilizing Descartes cell ontology signatures, revealing enrichment for stromal and extravillous trophoblast populations in relapsing tumors. (**C**) GSEA using canonical ontologies (Hallmark, KEGG, Reactome, WikiPathways), demonstrating massive enrichment of Epithelial-Mesenchymal Transition (EMT) and extracellular matrix (ECM) reorganization networks.ù

To contextualize the systemic biological consequences of this transcriptomic divergence, we performed Gene Set Enrichment Analysis (GSEA) contrasting relapsing versus non-relapsing M0 tumors. Rather than deploying canonical proliferative cascades, the functional landscape of metachronous relapse selectively amplified the extracellular remodeling and placental mimicry networks established in our TCGA analysis. Relapsing M0 tumors were defined by a massive, highly significant enrichment for Epithelial-Mesenchymal Transition (EMT), extracellular matrix (ECM) reorganization, and collagen assembly, indicating the active construction of a dense, desmoplastic niche [Fig. 2B]. Crucially, querying the Descartes cell ontology confirmed the physical composition of this remodeled niche [Fig. 2C]. The positive enrichment landscape was overwhelmingly dominated by generic stromal and vascular endothelial signatures across diverse tissue origins, physically validating a state of profound desmoplasia and active angiogenesis. However, embedded within this newly constructed vascular/stromal bed, we observed the striking retention of a highly specific oncofetal program, evidenced by the significant positive enrichment for “Extravillous Trophoblast”. Conversely, non-relapsing tumors retained signatures of aberrant epithelial differentiation (e.g., Keratinization, Cornified Envelope), suggesting that clinical relapse requires the complete abandonment of these static constraints in favor of a highly plastic, mesenchymal invasion. Operating within this heavily fortified extracellular matrix, relapsing M0 tumors exhibited a profound suppression of host adaptive immune signatures. We observed extreme negative enrichment across all T-cell activation, TCR signaling, and allograft rejection cascades, perfectly matched by a domain-wide depletion of lymphoid cell signatures in the Descartes ontology [Fig. 2C]. Together, these data confirm that the transcriptomic blueprint of metachronous relapse is defined by the coordinated execution of placental mimicry from within an immune-depleted, desmoplastic stroma. Given that *IGF2* emerged as the dominant transcriptional correlate of this extreme systemic phenotype, we focused our investigation on the specific dynamics of the 11p15.5 imprinted domain to dissect its underlying regulatory mechanics.

### Isoform analysis reveals an invariant oncofetal promoter architecture

Physiologically, *IGF2* is a highly complex locus subjected to strict spatio-temporal regulation governed by multiple distinct promoters (P1–P5), which dictate tissue-specific and developmental stage-specific expression [32]. Because dynamic promoter switching frequently drives phenotypic plasticity and aggressive state-shifts in cancer, we utilized proActiv to compute Relative Promoter Activity (RPA). Our objective was to determine whether the observed global upregulation in the relapsed group was driven by a qualitative structural shift or the quantitative amplification of a baseline fetal state.

The architectural landscape of *IGF2* transcripts was highly constrained and entirely monopolized by a dual-promoter system [Fig. 3A]. The P4 promoter, the dominant fetal driver, acted as the primary engine, accounting for the vast majority of transcript origins (∼65%), robustly followed by the downstream P5 promoter (∼30%), which is the human ortholog of the murine P0 promoter; while confined to the placenta in mice [33], in humans this promoter is also active in fetal skeletal muscle during development [34]. Notably, the adult-specific P1 promoter was entirely transcriptionally silent, while the fetal metabolic P3 promoter contributed only marginally (<5%). Crucially, analysis of the RPA matrix revealed no statistically significant differences in this structural hierarchy between the recurrent and non-recurrent groups. This invariant profile indicates that disease aggressiveness is fueled by a massive quantitative amplification of a pre-existing, truncal oncofetal program, rather than a topological reprogramming of internal promoter preference.

**Figure 3.**
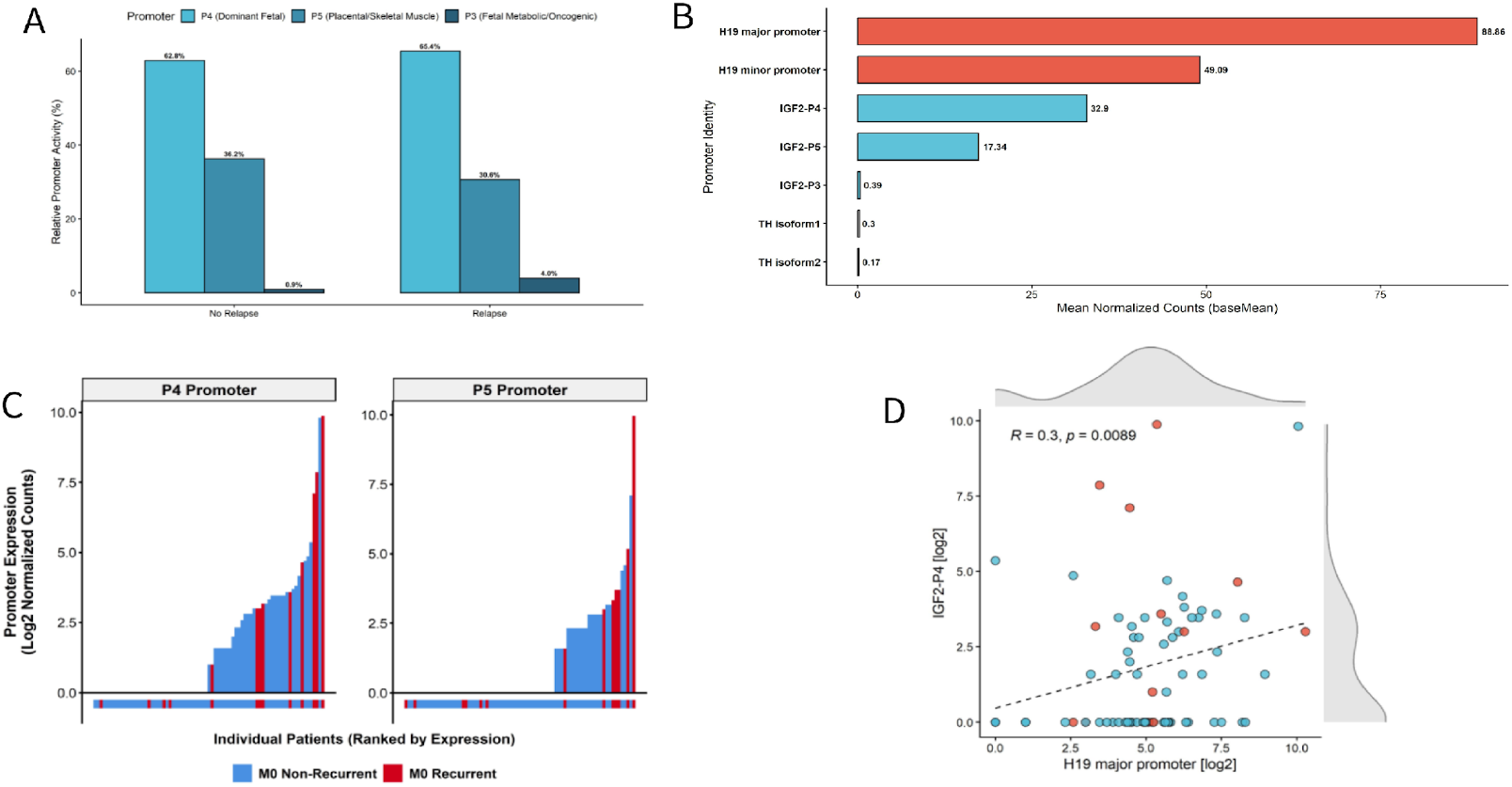
Transcriptional architecture and promoter-level dynamics of the IGF2/H19 imprinted locus in localized EOCRC. (**A)** Relative Promoter Activity (RPA) of *IGF2* transcripts stratified by clinical outcome (Relapse vs. No Relapse). The transcriptional landscape is dominated by the embryonic P4 and placental P5 promoters, with only marginal contribution from P3 and complete silencing of the adult P1 promoter. This architecture indicates a profound oncofetal transcriptional reversion that is established independently of metachronous relapse status. (**B**) Absolute transcriptional quantification (mean normalized counts) of the 11p15.5 domain regulatory elements. The data highlights massive co-activation of *H19* and fetal *IGF2* promoters, coupled with the strict topological silencing of the flanking boundary gene TH (Tyrosine Hydroxylase), supporting a localized epigenetic relaxation rather than broad chromosomal amplification. (**C)** Patient-level distribution of *IGF2* P4 and P5 promoter expression (log2 normalized counts). Individual M0 patients are ranked by expression magnitude, with color-coding indicating clinical outcome (Blue: Non-Recurrent; Red: Recurrent), demonstrating the continuum of oncofetal promoter activation across the cohort. (**D**) Spearman correlation analysis between the major *H19* promoter and the *IGF2* P4 promoter (*R = 0*.*3, p = 0*.*0089*). The significant positive correlation corroborates a coordinated, domain-wide epigenetic derepression mechanism overriding the canonical enhancer-competition model.

### Focal transcriptional derepression and coordinated *H19/IGF2* architecture

To map the physical extent of this deregulation, we expanded our absolute quantification to the individual promoters within the local 11p15.5 domain, encompassing *H19, INS/INS-IGF2*, and *TH*. We observed a massive transcriptional derepression specifically confined to the *IGF2* major promoters and the adjacent *H19* non-coding RNA, while the flanking genes *TH* and *INS-IGF2* remained strictly at physiological baseline (near-zero counts) across all patients [Figure 3B]. This strict spatial restriction broad chromosomal amplification as the driving mechanism, pointing instead to a focal epigenetic relaxation at the Imprinting Control Region (ICR1).

Interestingly, despite the massive global gene-level upregulation, promoter-level differential analysis (DESeq2) did not reach stringent statistical significance for *IGF2* individual promoters between the recurrent and non-recurrent groups. This uncoupling is driven by extreme intra-group dispersion: the cohort is characterized by a bimodal distribution comprising strict silencers and sporadic “hyper-expressor” tumors [Fig. 3C]. While hyper-expressors were enriched in the recurrent group, the existence of extreme hyper-expressors within the non-recurrent control group heavily inflated variance estimation. This bimodal distribution and incomplete clinical penetrance indicate that the hyperactivation of this embryonic growth factor represents a permissive transcriptomic state, but fundamentally insufficient on its own, to guarantee the execution of an aggressive systemic metastasis.

To determine if this focal deregulation represented a stochastic event or a structured domain-wide phenomenon, we analyzed the transcriptional coordination between the maternally-expressed *H19* and the paternally-expressed *IGF2*-P4 promoter. Scatterplot analysis confirmed a distinct bimodal transcriptional architecture, with a significant positive correlation between *H19* and *IGF2*-P4 absolute counts (Spearman R =, p = 0.0089) [Figure 3D]. This concomitant deregulation is highly suggestive of a coordinated epigenetic relaxation affecting the shared regulatory elements of the locus.

### Quantitative ASE Confirms Loss/Relaxation of Imprinting as a Truncal Oncofetal Event

To determine whether the extreme transcriptional bimodal architecture was driven by a relaxation of parent-of-origin expression control, we systematically interrogated a highly penetrant heterozygous single nucleotide polymorphism (SNP, *chr11:2,130,588*) in the *IGF2* gene across the M0 cohort [Fig. 4A, B]. Rather than a sporadic feature of extreme outliers, quantitative ASE revealed pervasive and statistically significant (*p < 0*.*05*, binomial test) biallelic IGF2 expression consistent with imprinting relaxation in 54 out of 77 M0 cases with adequate RNA-level evidence (70%). Strikingly, robust biallelic derepression (Loss/Relaxation of Imprinting, LOI) was not restricted to the “hyper-expressor” subset. Robust expression of the minority allele was consistently observed across the clinical spectrum, including baseline low-expressor and non-recurrent cases, converging on a median Variant Allele Frequency (VAF) of 0.37 [Fig. 4C]. Remarkably, the majority of these LOI-positive cases (50/54, 92.6%) were clinically confirmed as mismatch repair proficient (MSS), with only a marginal fraction (n=4) exhibiting distinct MMR deficiency profiles. The identification of this persistent biallelic signature—irrespective of the final transcriptional magnitude or clinical relapse status— fundamentally redefines LOI at the *IGF2* locus as an early, truncal event in EOCRC tumorigenesis. It may provide an oncofetal permissive state, while the subsequent quantitative amplification and additional adaptive programs appears necessary for systemic failure.

**Figure 4:**
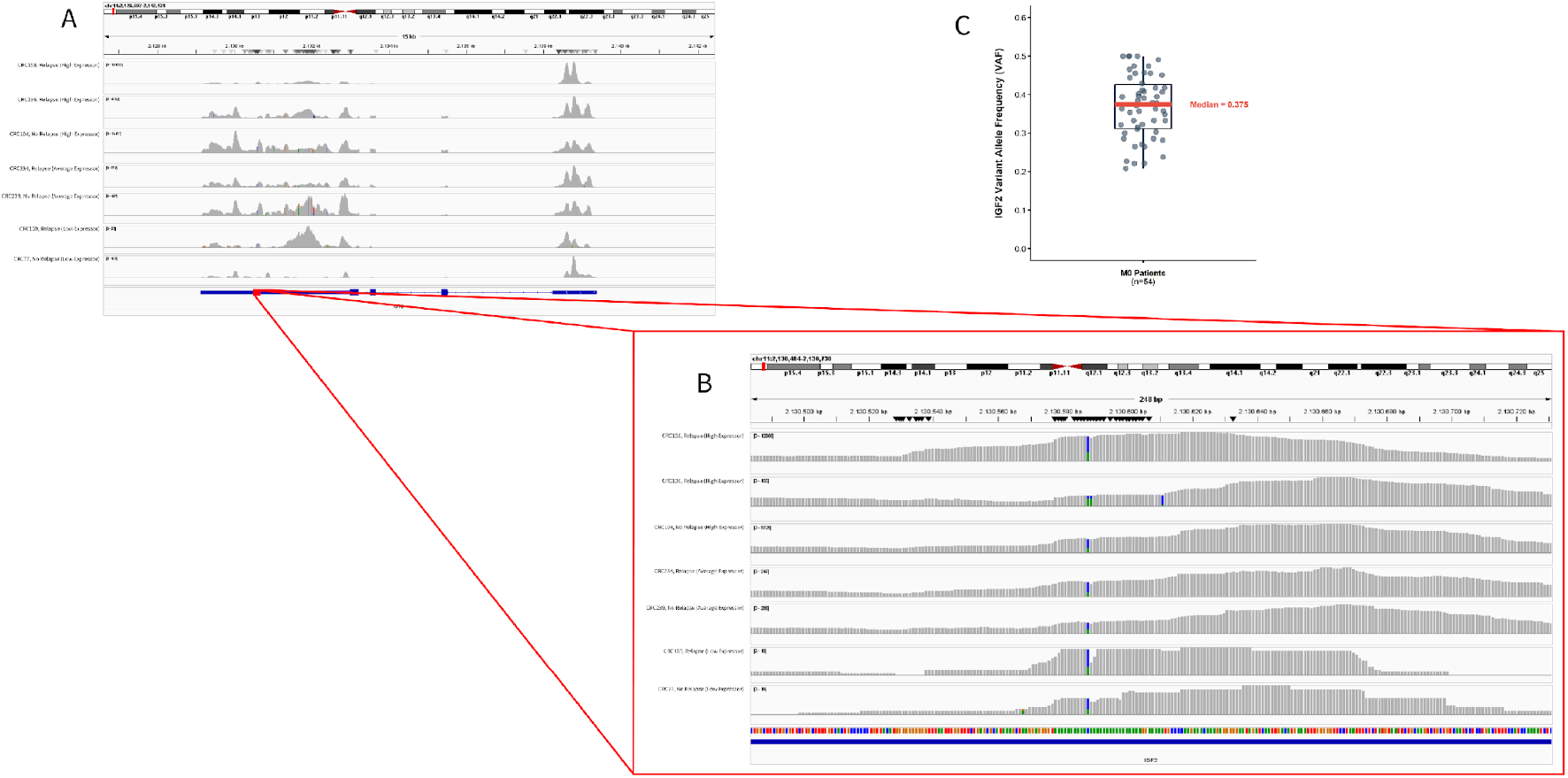
Systematic Allele-Specific Expression (ASE) profiling confirms structural Loss/Relaxation of Imprinting (LOI) at the IGF2 locus. (**A**) Integrative Genomics Viewer (IGV) visualization of RNA-seq read alignments across the extended 11p15.5 imprinted domain for representative EOCRC index cases exhibiting different target expression. (**B**) High-resolution transcriptomic phasing at the highly penetrant heterozygous SNP chr11:2,130,588 located in the coding exons of *IGF2* gene. The concurrent presence of distinct sequence alleles (reference and alternate) within the read pileups indicates active biallelic transcription, consistent with relaxation of physiological monoallelic silencing. (**C**) Quantitative assessment of transcriptomic Variant Allele Frequency (VAF) across the informative M0 cohort (n=54). The cohort exhibits pervasive biallelic transcription with a median VAF of 0.37, an allelic frequency mathematically consistent with robust biallelic tumor expression diluted by physiological monoallelic stromal admixture, indicatingIGF2 imprinting relaxation as a pervasive truncal event in EOCRC.

### Bootstrap-validated LASSO regression defines a multi-class adaptive survival module

As demonstrated by our cohort dispersion, the biallelic hyperactivation of *IGF2* establishes a potent oncofetal baseline, yet it remains functionally insufficient on its own to execute a systemic relapse. We hypothesized that to survive the extreme microenvironmental stress of dissemination, *IGF2*-driven clones must recruit a secondary transcriptomic network governing cellular plasticity and adaptive tolerance. To objectively distill this complex adaptive landscape without *a priori* biological bias, we subjected the nominal differential expression space (univariate p < 0.05) to LASSO-penalized Cox regression. Using bootstrap resampling (n=1,000 iterations) to estimate feature stability and reduce overfitting risk, this unbiased regularization identified an internally derived 5-gene recurrence risk module: the *IGF2* engine, *UTS2B*, a potent endogenous vasoactive peptide; *ATG10*.*IT1*, an intronic transcript housed within the core autophagic *ATG10* locus; *UBE2SP2*, a transcribed pseudogene belonging to the ubiquitin-conjugating enzyme family; and *LINC01730*, a long non-coding RNA [Fig. 5A]. All five genes were retained in ≥30% of iterations, with *IGF2* present in >70% of resamples. Crucially, the predictive performance of this module was rigorously validated across the resampling framework. Time-dependent Receiver Operating Characteristic (ROC) analysis confirmed robust internal consistency, yielding high mean Area Under the Curve (AUC) profiles at 12, 24, and 36 months [Fig. 5B]. Translating this continuous transcriptomic risk into a dichotomized clinical threshold, Kaplan-Meier survival analysis demonstrated that the acquisition of this composite signature profoundly accelerates metachronous relapse, effectively separating aggressive systemic trajectories from localized, disease-free persistence [Fig. 5C, p < 0.0001].

**Figure 5.**
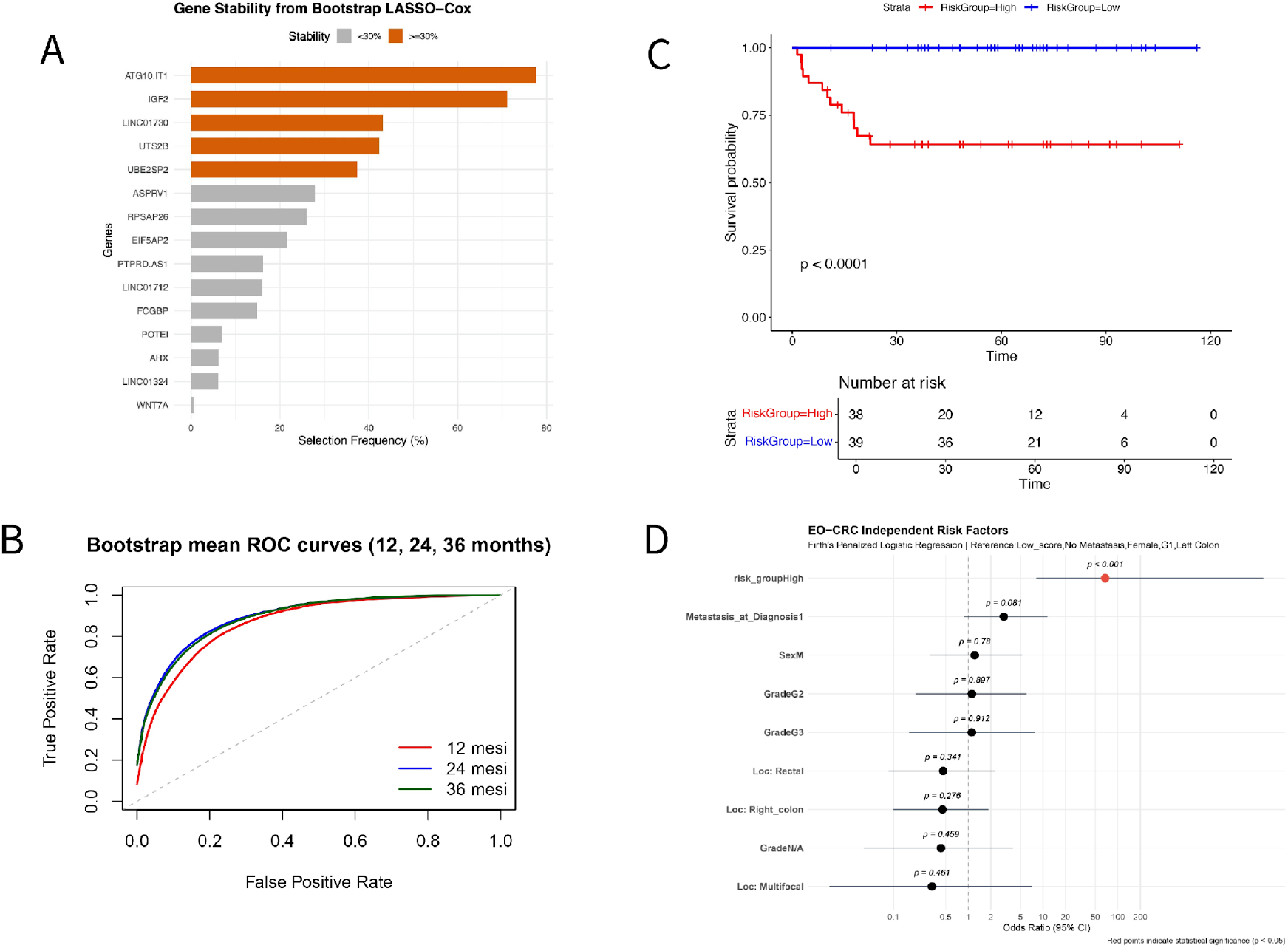
Identification and clinical validation of a 5-gene recurrence-risk module. (**A**) Gene selection stability plot from bootstrap-resampled (n=1,000) LASSO-Cox regression, identifying a robust 5-gene module (*ATG10*.*IT1, IGF2, LINC01730, UTS2B, UBE2SP2*). (**B**) Bootstrap mean time-dependent ROC curves evaluating the predictive accuracy of the signature at 12, 24, and 36 months. (**C**) Kaplan-Meier survival analysis stratifying patients by the composite 5-gene risk score (p < 0.0001). (**D**) Forest plot of multivariable Firth’s Penalized Logistic Regression in a curative-intent pan-cohort (n=96), confirming the high-risk transcriptomic group as independently associated with metastatic events (p < 0.001) when adjusted for standard clinical variables.

### Multivariable Analysis Indicates Prognostic Independence from Pathological Covariates

To further validate the prognostic performance of the 5-gene module under conditions of extreme biological aggressiveness, we tested its prognostic power in a pure, synchronous metastatic (M1) cohort that had achieved a transient, macroscopically disease-free state following curative-intent metastasectomy (n=19). Even within this uniformly advanced background where standard clinical staging is fundamentally uninformative, the signature accurately identified the patients (n=15) who subsequently experienced a secondary systemic relapse, proving its capacity to faithfully capture residual disease capacity even within this limited cohort (p = 0.0026).

To then establish whether this multi-class module is associated with outcome independently of late-stage presentation, we scaled our analysis to a combined, strictly defined curative-intent pan-cohort (M0 and fully resected M1 patients, n=96) utilizing Firth’s Penalized Logistic Regression. The multivariable analysis of 96 patients revealed that the high-risk group was the only variable significantly associated with metastatic events (OR = 67.71; 95% CI: 8.07–8870.40; p < 0.001) (Figure 5 D). While patients presenting metastasis at diagnosis showed a trend toward higher odds of the event (OR = 2.98; 95% CI: 0.88–11.48), this did not reach statistical significance (p = 0.081). Other clinical variables, including sex, tumor grade, and anatomical location (rectal, right colon, or multifocal), showed no significant association with the outcome (p>0.05 for all). The overall model fit was highly significant (Likelihood Ratio Test p<0.001).

## Discussion

The identification of *IGF2* as the dominant transcriptional correlate of metachronous relapse within our localized M0 cohort positions the 11p15.5 imprinted domain as a central vulnerability axis in early-onset disease. *IGF2* LOI is an established epigenetic event in colorectal carcinogenesis, detectable as a constitutional field defect in normal colonic mucosa of approximately 30% of CRC patients and even in peripheral blood lymphocytes, where it confers an adjusted odds ratio of 21.7 for concurrent CRC diagnosis [35]. Functional modelling confirmed that *Igf2* LOI in mice is causally sufficient to double intestinal tumor formation and to shift the normal colonic epithelium toward a progenitor-enriched state [36].

Our data advance this framework by providing a quantitative, promoter-resolved characterization of LOI specifically within an early-onset cohort. Our ASE analysis revealed pervasive biallelic transcription with sub-clonal allelic frequencies mathematically consistent with stromal admixture. Remarkably, the overwhelming majority of these LOI-positive cases were mismatch repair proficient (MSS), with only a marginal fraction exhibiting MMR deficiency profiles. This distribution uncouples *IGF2* epigenetic collapse from the CpG island methylator phenotype (CIMP) associated with MSI tumors. Instead, it anchors this focal epigenetic relaxation as a candidate truncal feature of genomically stable early-onset disease, mirroring the oncofetal baseline established in our TCGA analysis.

Critically, the topological restriction of this deregulation argues against large-scale chromosomal amplification, pointing instead to a focal epigenetic relaxation at the shared ICR1 governing the *IGF2/H19* domain, consistent with the hypomethylation-based LOI mechanism previously characterised in CRC [37; 38]. The significant positive correlation between *H19* and *IGF2*-P4 absolute counts further supports coordinated domain-wide epigenetic relaxation. This interpretation is consistent with the proposal that the epigenetic alteration responsible for abnormal *IGF2* imprinting in CRC patients represents an embryogenic event predating tumor formation [39], and is reinforced by the independent association of *IGF2* DMR0 hypomethylation with significantly worse overall survival [40]. Our data extend this prognostic association by demonstrating that within the early-onset context, it is not the LOI event itself but the degree of quantitative amplification superimposed upon it that stratifies metachronous relapse risk.

Within this framework, the promoter-level architecture we resolved in EOCRC introduces a critical distinction from previous CRC literature. The available evidence on *IGF2* promoter dynamics in colorectal tumorigenesis rests primarily on in vitro data: Singh and colleagues demonstrated co-activation of P3 and P4 promoters in proliferating Caco-2 colon cancer cells, with both declining upon differentiation and P1 remaining entirely inactive [41]. Based on this and related functional evidence — including the identification of *PLAG1* as a transcription factor capable of driving *IGF2* expression through P3 binding in a cell-type specific manner [42] — subsequent reviews have described P3 reactivation as the predominant fetal promoter mechanism in CRC [43]. However, no study has applied high-resolution, RNA-seq-based promoter quantification to primary CRC tumor tissue to directly test this inference. Our proActiv analysis of primary EOCRC tumors reveals a markedly distinct architecture: near-complete dominance of P4 (the primary embryonic driver) and the P5 placental promoter, with P3 contributing only marginally, and the adult-specific P1 remaining entirely silent. This departure from the cell-line-inferred P3-dominant model is biologically coherent in the specific context of early-onset disease: rather than recapitulating the proliferative-phase dynamics of a cycling colon cancer cell line, EOCRC appears to recruit a more archaic, embryonically anchored transcriptional program. The dominance of P4 and P5 is fully consistent with the trophoblastic mimicry identity established in our TCGA exploratory analysis, and reinforces the ontogenetically distinct nature of EOCRC.

However, establishing the primary niche is fundamentally distinct from surviving systemic dissemination. This functional insufficiency of *IGF2* LOI alone to guarantee metastatic execution provides the biological rationale for the multi-class adaptive survival module identified through our bootstrap-validated LASSO-Cox framework. This module couples the fundamental IGF2 engine with the vasoactive peptide *UTS2B*—a plausible counterweight to hemodynamic stress during peripheral dissemination —and a specific non-coding framework (*UBE2SP2, LINC01730*, and *ATG10*.*IT1*).

Crucially, the retention of elements like the intronic transcript *ATG10*.*IT1* and *LINC01730* must be interpreted through the lens of the profound RNA splicing hyperactivity observed in EOCRC, as demonstrated in our exploratory TCGA analysis. This hyperactive RNA-processing infrastructure plausibly licenses the stable expression of intronic and non-coding transcripts that would remain below detection thresholds in transcriptionally quiescent tumor contexts. Rather than functioning as isolated, cell-autonomous mechanistic drivers, these elements serve collectively as molecular sentinels, acting as surrogate biomarkers reflecting the global epigenetic fluidity and transcriptional plasticity required for disseminated clone survival.

Notably, while prior studies link *IGF2* expression [43,44] and protein positivity [45] with worse clinical outcomes across unselected CRC cohorts, our multivariate framework indicates that, within the EOCRC context, the combined transcriptomic module adds critical prognostic information beyond standard staging variables.

Several limitations of this study warrant explicit acknowledgment. First, while our ASE analysis identified pervasive biallelic transcription, definitive exclusion of somatic heterozygosity at the interrogated locus formally requires paired germline DNA sequencing, which was unavailable. Furthermore, while TCGA data firmly establish the oncofetal baseline, orthogonal transcriptomic profiling of strictly matched late-onset CRC cohorts will be required to confirm that this developmental mimicry is a hallmark of early-onset disease. Second, while bootstrap resampling provides rigorous internal stability estimates, retrospective *in silico* validation of the 5-gene signature was strictly precluded by the structural limitations of currently available public transcriptomic repositories. Existing public CRC datasets consistently fail to meet the overlapping prerequisites for this specific validation: they typically lack sufficient statistical power for early-onset cohorts, do not provide granular Recurrence-Free Survival (RFS) annotations for localized disease, or rely on legacy microarray platforms fundamentally incapable of capturing the specific low-abundance non-coding and intronic transcripts (e.g., *ATG10*.*IT1, LINC01730*) integral to our survival module. Consequently, definitive confirmation of the signature’s prognostic utility must rely on independent validation within a dedicated, prospective cohort utilizing standardized RNA-seq profiling, where the inherent sensitivity of non-coding elements to pre-analytical RNA quality variance can be rigorously controlled.

In summary, our findings provide a working model that reframe the systemic aggressiveness of EOCRC as the product of a dual-hit architecture: the focal deregulation of the IGF2/H19 domain as a candidate oncofetal permissive event, and the subsequent acquisition of a multi-class adaptive survival module as recurrence-associated transcriptional program. This framework resolves the paradox of widespread *IGF2* LOI co-existing with heterogeneous outcomes. The primary tumor-level promoter resolution establishes the EOCRC epigenetic landscape as more developmentally archaic than previously inferred from cell-line models, offering a biologically grounded basis for the molecular risk stratification of EOCRC patients outside the boundaries of anatomical staging.

## Methods

### Ethics Approval and Patient Enrollment

Patient enrollment was conducted after obtaining approval from the appropriate Ethics Committee (Protocol ID: DOC2RES 18395).

### RNA extraction and library preparation for RNA sequencing

RNA was isolated from 5 µm formalin-fixed paraffin-embedded (FFPE) tissue sections using the EZ2 RNA FFPE Kit (Qiagen, Milan, Italy), following the manufacturer’s protocol. RNA concentration was quantified using the Qubit Fluorometric System, while RNA quality was assessed using the Agilent TapeStation. Sequencing libraries were generated using the Illumina Stranded Total RNA Prep Ligation with Ribo-Zero Plus according to the manufacturer’s recommendations. Paired-end sequencing (2 × 101 bp) was performed on the NovaSeq 6000 platform (Illumina, San Diego, CA, USA).

### RNA-seq Data Processing and Quality Control

Raw sequencing reads in FASTQ format were subjected to quality assessment using FastQC (v0.11.9) to evaluate per-base quality scores, GC content, sequence duplication levels, and adapter contamination. Potential cross-species contamination was screened using FastQ Screen (v0.14.1), which aligns reads against multiple reference genomes in parallel.

Primary bioinformatic analysis was executed through a custom pipeline developed in R. High-quality reads were aligned to the ENSEMBL reference genome using the STAR alignment algorithm, generating corresponding BAM and BAI files. Subsequently, transcripts and gene-level quantification were performed using RSEM to obtain raw read counts. Post-alignment quality control and assessment of mapping statistics for each BAM file were systematically conducted using QualiMap. Gene-level count matrices were imported into RStudio (R v4.3.1) for downstream analysis. Genes with fewer than 10 counts in at least 5% of samples were excluded to remove low-abundance features. Technical replicates were collapsed into a single representative profile per patient using the collapseReplicates function from the DESeq2 package.

### Public Data Acquisition and Cohort Stratification

Transcriptomic profiles (HTSeq-Counts) and corresponding clinical metadata from the TCGA Colorectal Adenocarcinoma (COADREAD) cohort were retrieved from the GDC Data Portal (https://portal.gdc.cancer.gov/) using the TCGAbiolinks R package, ensuring access to the most recent data alignment against the GRCh38 (hg38) human reference genome. To isolate the intrinsic, baseline molecular architecture of the primary tumor and eliminate confounding effects related to therapeutic interventions, we strictly restricted the analysis to treatment-naive patients. To further minimize biological noise and isolate age-specific biological drivers, we implemented a stringent “hard-filtering” pipeline, retaining only primary tumor samples with a confirmed histological diagnosis of of Adenocarcinoma (Not Otherwise Specified; NOS), excluding variant histologies (e.g., mucinous or signet-ring cell carcinomas), non-primary tissues, and cases with incomplete pathologic staging. To ensure a biologically homogeneous background, we integrated molecular subtyping data from the TCGA Pan-Cancer Atlas (2018) [46]. The cohort was filtered to include only Microsatellite Stable/Chromosomal Unstable (MSS/CIN) tumors, formally excluding MSI-High, POLE-mutated, and hypermutated non-MSI phenotypes. Following the removal of low-count transcripts (defined as >10 raw counts in at least 25% of the smallest experimental group), the final curated dataset was stratified into three chronological cohorts: n= 39 Early-Onset (EO; <50 years) patients, n=144 Average-Onset (AO; 50–70 years) patients, and n=129 Late-Onset (LO; >70 years).

### TCGA-Differential Gene Expression Analysis (DGEA)

Differential Gene Expression Analysis (DGEA) was performed using the DESeq2 R package (v1.4x). To account for potential clinical confounders, a Generalized Linear Model (GLM) was parameterized with a multi-factor design formula, that explicitly adjusted for gender, AJCC pathologic stage, and anatomical location (Right Colon, Left Colon, and Rectum) to isolate age-dependent transcriptional shifts. Dispersion and log2 Fold Change (*LFC*) estimates were calculated for the entire cohort to maximize statistical power. Specific contrasts (EO vs. LO and EO vs. AO) were extracted using the Wald test. To refine effect size estimates and minimize the impact of high-dispersion outliers, *LFC* values were corrected using the apeglm (Approximate Posterior Estimation for GLM) shrinkage estimator [47]. Significant differentially expressed genes (DEGs) were defined by a False Discovery Rate (FDR-adjusted p-value) < 0.05 and an absolute |LFC| > 0.5.

### TCGA-Gene Set Enrichment Analysis (GSEA) and Ontogenetic Mapping

Functional enrichment was assessed via Gene Set Enrichment Analysis (GSEA) using the clusterProfiler and ReactomePA packages. All expressed genes were pre-ranked by multiplying the sign of the log2 Fold Change (+1 for upregulation, -1 for downregulation) by the corresponding -log10(p-value) of each gene. This ranking prioritized genes demonstrating both high magnitude of change and robust statistical significance. The ranked lists were interrogated against curated molecular signatures from MSigDB (Hallmarks, Gene Ontology, KEGG, Descartes cell type and Reactome). The GSEA algorithm was executed with 1,000 permutations, with a minimum gene set size of 15 and a maximum of 500. Statistical significance for pathway enrichment was defined by p-value < 0.05 and a Normalized Enrichment Score (NES) indicating the direction of the effect. Multiple testing was controlled via FDR, with a threshold of < 0.1 for detailed biological interpretation.

### Internal EOCRC cohort-Differential Gene Expression and Pathway Analysis

Normalization and differential expression analysis were performed using DESeq2. Variance-stabilizing transformation (VST) was applied to normalized counts for visualization and exploratory analyses. Residual batch effects were corrected using the removeBatchEffect function from the limma package.

Differentially expressed genes (DEGs) were defined by an adjusted p-value < 0.05 and an absolute log_2_ fold change ≥ 1.5. Gene Set Enrichment Analysis (GSEA) was performed on a pre-ranked gene list ordered by the DESeq2 Wald statistic. The following MSigDB gene set collections were interrogated: Hallmarks of Cancer, KEGG, Reactome, Descartes Cell Types and Tissue and WikiPathways.

### Promoter-Level Activity Quantification and Isoform Switching Analysis

To investigate transcriptional dynamics and differential promoter usage at the 11p15.5 locus, absolute promoter activity was quantified from spliced RNA-seq alignments using the proActiv framework, based on the GENCODE v47 (GRCh38) reference annotation. Differential promoter activity between recurrent and non-recurrent tumors was evaluated using the DESeq2 R package. Promoters with fewer than 10 total counts across the cohort were excluded prior to downstream modeling.

To prevent normalization artifacts arising from the targeted analysis of a highly deregulated topological associated domain (TAD), library size factors were estimated from the complete genome-wide promoter count matrix (>70,000 promoters). To address the inherent sparsity of transcript-level quantifications, global size factors were computed utilizing a zero-tolerant estimator (type = “poscounts” in DESeq2), which calculates the geometric mean strictly from positive counts. These globally anchored, sparsity-corrected size factors were subsequently applied to the targeted 11p15.5 DESeq2 model (encompassing *IGF2, H19, TH, INS*, and *INS-IGF2*). Logarithmic fold-change estimates were shrunken using the apeglm method to stabilize the variance of low-count promoters [47].

To evaluate structural shifts in transcript composition, Relative Promoter Activity (RPA) was calculated as the fractional contribution of each specific promoter to the total transcriptional output of its parent gene. This compositional metric was strictly employed to delineate internal alternative promoter usage and isoform switching at the IGF2 locus, specifically evaluating the balance among the P3, P4, and P5 promoters. Statistical differences in RPA between clinical outcomes were assessed using the Wilcoxon rank-sum test, with p-values adjusted via the Benjamini-Hochberg False Discovery Rate (FDR) method to account for multiple comparisons.

### Allele-Specific Expression (ASE) Profiling

To mechanistically validate the Loss of Imprinting (LOI) predicted by the bulk transcriptomic deregulation of the 11p15.5 locus, we performed a systematic, read-level Allele-Specific Expression (ASE) analysis. Binary Alignment Map (BAM) files—aligned against the GRCh38 reference genome—were systematically interrogated to assess the transcriptional status of a highly penetrant heterozygous single nucleotide polymorphism (SNP, chr11:2,130,588) mapped to the *IGF2* transcribed region. Across the localized EOCRC cohort (n=77), 54 showed sufficient RNA-level allelic evidence at the interrogated coordinate to support biallelic expression.

Allelic expression status was quantitatively evaluated by extracting the allele-specific RNA read counts across the targeted coordinate to calculate the transcriptomic Variant Allele Frequency (VAF). Maintenance of physiological imprinting (monoallelic expression) was strictly defined by a VAF approaching 0 or 1, representing the exclusive transcription of a single parental allele. Conversely, the epigenetic relaxation of imprinting (biallelic expression) was established by intermediate, sub-clonal VAFs, indicating the simultaneous transcription of both the reference and alternate alleles. This transcriptomic VAF profiling provided a direct, quantitative molecular readout of the active recruitment of both parental chromosomes by the RNA Polymerase II machinery. While paired germline DNA was unavailable to formally confirm constitutional heterozygosity across all samples, the detection of transcriptomic biallelic expression indicates localized epigenetic relaxation (LOI) in the informative subset.

### Prognostic Signature Identification and Internal Validation

Relapse-free survival (RFS) analysis was conducted using the survival R package. Candidate prognostic genes were first identified through univariate Cox proportional hazards regression applied to all DEGs, retaining genes with p < 0.05. Selected candidates were then subjected to LASSO-penalized Cox regression using the glmnet package. The optimal regularization parameter (λ) was determined by 10-fold cross-validation, and genes with non-zero coefficients at λ.min were retained for signature construction.

To assess the stability and reproducibility of the prognostic signature, a bootstrap-based variable selection framework was implemented across 1,000 iterations. In each iteration, a bootstrap sample was generated by resampling patients with replacement. Within each resampled cohort: (i) candidate genes were pre-screened by univariate Cox regression (p < 0.05); (ii) LASSO-penalized Cox regression was applied to the pre-screened gene set; and (iii) model performance was evaluated using the concordance index (C-index) and time-dependent AUC at 12, 24, and 36 months. Genes selected in ≥ 30% of bootstrap iterations were retained in the final signature.

An individual risk score for each patient j was computed as a weighted linear combination of VST-normalized expression values:

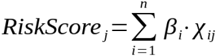

where *n* denotes the number of genes in the final signature, *β_i* is the Cox regression coefficient for gene *i*, and *x_ij* is the VST-normalized expression value of gene *i* in patient *j*. Risk scores were subsequently scaled and patients were dichotomized into “High” and “Low” risk groups using the median risk score as the stratification threshold.

### Multivariate Analysis and Independence Testing

To identify clinical predictors of metastatic events (EVENT_MET), a multivariable logistic regression model was employed. To account for potential issues of small sample size and quasi-separation among predictors, we used Firth’s Penalized Likelihood Logistic Regression (via the logistf R package). The model included risk group, metastasis at diagnosis, sex, tumor grade, and tumor location as independent variables. Confidence intervals (95% CI) and p-values were calculated using the profile penalized log-likelihood method, which provides more reliable estimates than standard Wald tests in the presence of rare events.

## Acknowledgements

Funded by PNRR-MAD-2022-12376835 Funded by the European Union - Next Generation EU - M6C2 - Investment 2.1 Dissecting the biology of early-onset colorectal cancer C53C22001150007. We thank the Genomics GSTeP Facility of Fondazione Policlinico Universitario Agostino Gemelli IRCCS for technical support in RNA sequencing and genomic data generation. We also thank Cristina Vacca for her valuable support as study coordinator in the organization and management of the EOCRC study.

## Data availability

The TCGA data used in this study are publicly available through the Genomic Data Commons. The internal EOCRC RNA-seq dataset contains patient-derived genomic and clinical information and is not publicly available at the preprint stage. De-identified data supporting the findings of this study will be made available upon reasonable request and/or deposited in a controlled-access repository upon publication in a peer-reviewed journal, in accordance with institutional, ethical, and data-protection requirements.

## Code availability

Custom analysis code is not publicly available at the preprint stage. Scripts required to reproduce the main analyses will be made available upon reasonable request and/or released upon publication in a peer-reviewed journal.

## Ethics approval and consent to participate

This retrospective study was conducted in accordance with the Declaration of Helsinki and was approved by the Comitato Etico Territoriale Lazio Area 3 – Roma / CET Lazio Area 3, Fondazione Policlinico Universitario Agostino Gemelli IRCCS–Università Cattolica del Sacro Cuore, Rome, Italy (Protocol code: EO-CRC; CINECA ID: 5527; last favorable opinion issued on 19 March 2026). The requirement for written informed consent was waived by the Ethics Committee due to the retrospective nature of the study and the use of pseudonymized archival samples and clinical data, in accordance with applicable regulations.

