## Supplementary material for "An IGF2-anchored oncofetal state and relapse-associated transcriptomic module define systemic progression risk in early-onset colorectal cancer": Table 1 and supplementary Figures

| Characteristic | Total Cohort (n=96) | M0 Non-Relapse (n=64) | M0 Relapse (n=13) | M1 (n=19) |
|---|---|---|---|---|
| **Demographics** | | | | |
| Ag, median (range) | 44 | 44 | 43 | 44 |
| Sex (Male/Female) | M= 45 F=51 | M= 32 F=32 | M= 8 F=5 | M= 5 F=14 |
| **Clinicopathological Features** | | | | |
| Primary Tumor Location | Left=35 Right=30 Rectum=18 Rectosigmoid junction=5 Transverse=6 Synchronous multifocal=2 | Left=19 Right=24 Rectum=14 Rectosigmoid junction=3 Transverse=4 Synchronous multifocal=0 | Left=6 Right=3 Rectum=2 Rectosigmoid junction=1 Transverse=0 Synchronous multifocal=1 | Left=10 Right=3 Rectum=2 Rectosigmoid junction=1 Transverse=2 Synchronous multifocal=1 |
| pT Stage | T1-T2=18 T3-T4= 66 other (pT0-is N/A)=12 | T1-T2=15 T3-T4= 40 other (pT0-is N/A)=9 | T1-T2=0 T3-T4= 13 other (pT0-is N/A)=0 | T2=3 T3-T4= 13 other (N/A)=3 |
| pN Stage | pN0=39 pN+=47 N/A=10 | pN0=33 pN+=24 N/A=7 | pN0=2 pN+=11 N/A=0 | pN0=4 pN+=12 N/A=3 |
| Tumor Grade | G1-G2= 59 G3=26 N/A=11 | G1-G2= 40 G3=18 N/A=6 | G1-G2= 7 G3=5 N/A=1 | G1-G2= 12 G3=3 N/A=4 |
| Lymphovascular Invasion | Absent=64 Present= 32 | Absent=48 Present= 16 | Absent=4 Present= 9 | Absent=12 Present= 7 |
| Perineural Invasion | Absent= 65 Present=31 | Absent= 48 Present=16 | Absent= 6 Present=7 | Absent= 11 Present=8 |
| MMR Status (pMMR/dMMR) | pMMR= 58 dMMR=8 N/A=30 | pMMR=30 dMMR=8 N/A=26 | pMMR=12 dMMR=0 N/A=1 | pMMR=15 dMMR=1 N/A=3 |
| **Treatment** | | | | |
| Neoadjuvant/Adjuvant Chemotherapy | N/A= 24 YES=60 NO=12 | N/A= 12 YES=30 N=22 | N/A= 0 YES=13 N=0 | N/A= 2 YES=17 N=0 |
| **Clinical Outcome** | | | | |
| Follow-up | n=96 | n=64 | n=13 | n=19 |
| DFS (%) | 75% | 100% | 31% | 21% |

**Supplementary Table 1. Baseline demographic, clinicopathological, treatment, and clinical outcome characteristics of the study cohort stratified according to metastatic status and relapse occurrence.** Abbreviations: M0, non-metastatic disease at diagnosis; M1, metastatic disease at diagnosis; LVI, lymphovascular invasion; PNI, perineural invasion; MMR, mismatch repair; pMMR, proficient mismatch repair; dMMR, deficient mismatch repair, DFS, disease-free survival.

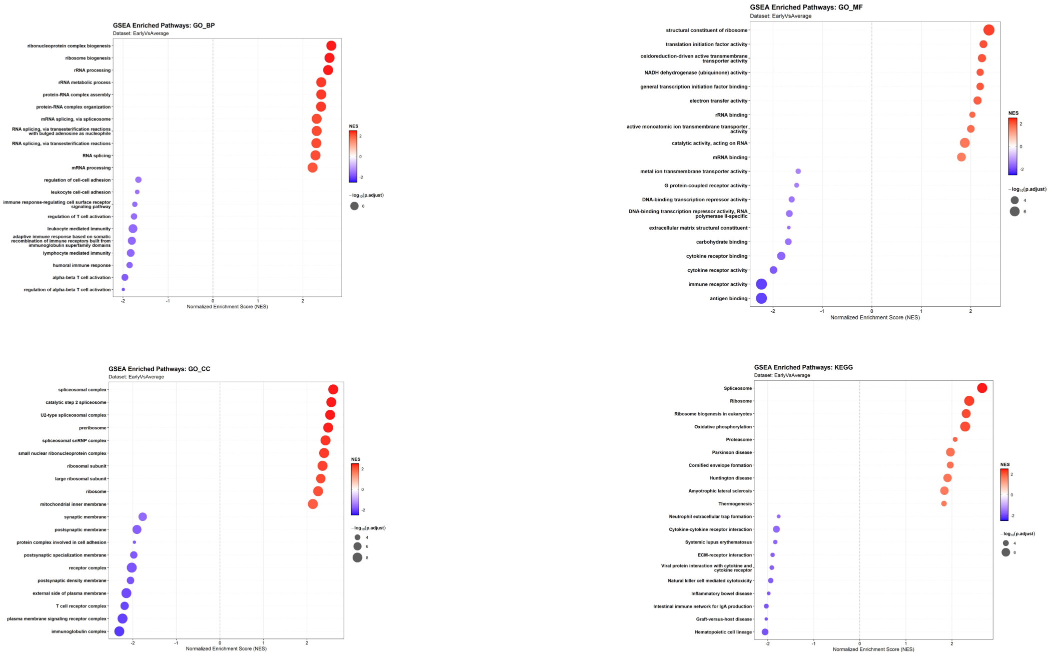


**Figure S1. Gene Set Enrichment Analysis (GSEA) for the Early versus Average comparison.** Dot plots depict functional enrichment results across Gene Ontology (GO) Biological Process (top-left), GO Molecular Function (top-right), GO Cellular Component (bottom-left), and KEGG (bottom-right) ontologies

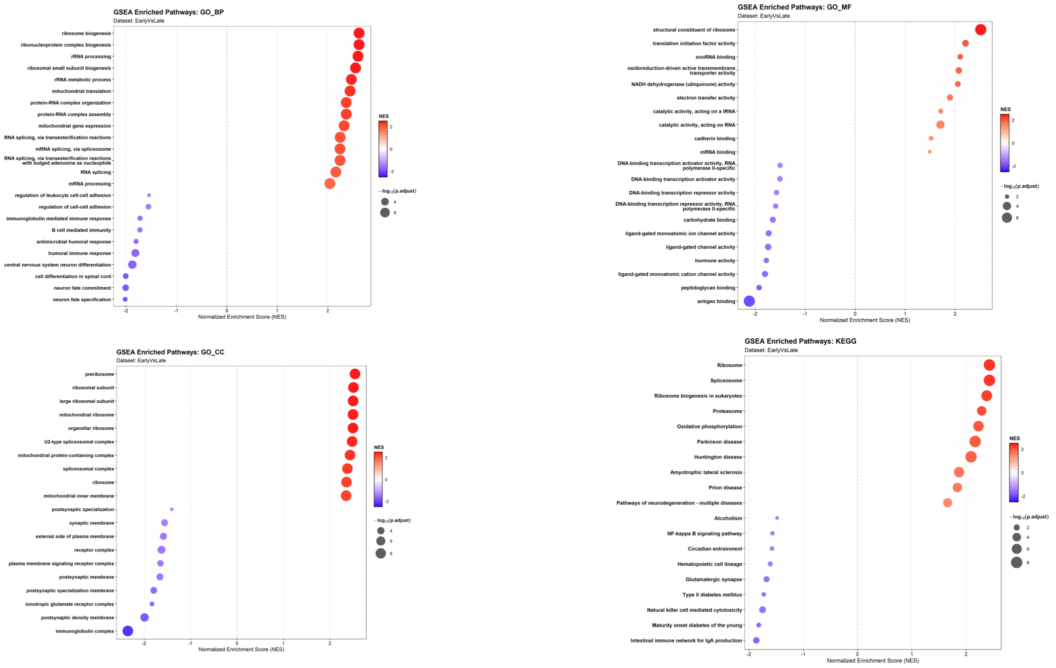


**Figure S2. Gene Set Enrichment Analysis (GSEA) for the Early versus Late comparison.** Dot plots depict functional enrichment results across Gene Ontology (GO) Biological Process (top-left), GO Molecular Function (top-right), GO Cellular Component (bottom-left), and KEGG (bottom-right) ontologies.
